# Spatiotemporal dynamics of RNA virus diversity in a phyllosphere microbial community

**DOI:** 10.1101/772475

**Authors:** Lisa M. Bono, Richard J. Orton, Elena L. Peredo, Hilary G. Morrison, Mark Sistrom, Sheri Simmons, Paul E. Turner

## Abstract

Although metagenomics reveals that natural virus communities harbor vast genetic diversity, the spatiotemporal dynamics of viral diversity in the wild are seldom tested, especially across small geographic scales. This problem is usefully examined in the above-ground phyllosphere, because terrestrial plants are frequently infected by taxonomically-diverse RNA viruses, whose elevated mutation rates generate abundant allele diversity. Here, we studied the problem by comparative analysis of RNA virus samples over time from three spatially-separated patches of a common perennial legume, white clover (*Trifolium repens* L.), growing in a grassy lawn in Woods Hole Village (Falmouth, MA, USA). We predicted that clover samples would show similarly high levels of virus species (alpha) diversity across space, but differing among-patch diversity of non-dominant virus taxa over time (4 samples spanning 6 weeks). Results showed that recognizable alpha diversity in clover patches was consistently dominated by RNA virus family *Alphaflexiviridae* across space, but that all patches showed inconsistent spatiotemporal presence of a diversity of minority virus families. Also, we observed that white clover mosaic virus (WClMV) dominated all patches across space and time. The high coverage of WClMV fostered an haplotype analysis, which revealed that two strains of the virus consistently infected clover plants during the 6-week period.

## Introduction

Viruses numerically dominate biological entities on Earth and may represent the majority of its genetic diversity as well [1, 2]. Viruses can alter microbial community structure [3] and colonization [4] and impact food webs. Their effects also scale up, driving global geochemical cycles [5, 6]. However, virus sampling across different natural environments is uneven. Marine systems are perhaps the best studied to date [7], making it crucial to examine non-marine and terrestrial communities (but see [4, 8]). Understanding the viral communities of the phyllosphere (above-ground terrestrial plant biomass) has important implications for food safety, *e.g*. outbreaks of food-borne human pathogens, such as hepatitis A virus and norovirus, have been associated with fresh produce [9]. Furthermore, the broadscale ecological and evolutionary implications of plant viruses is largely unknown, even though viral infections have been detected in a variety of wild plants [10]. Even a seemingly basic question about the nature of viral communities associated with plants remains unknown: will the same viruses be observed today as they will tomorrow, the next week, *et cetera*?

Plant viruses face unique challenges when infecting their hosts, which can potentially drive the ecological and evolutionary dynamics of their microbial communities. The plant cell wall is particularly difficult to breach; viruses can avoid this issue either through vertical transmission, or by relying on passive, mechanical damage to plant cell walls, most notably via feeding by phytophagous insects but also via animal grazing and human activities such as lawn mowing. This differs from phage infection of bacterial cells and viral infection of animal cells, where viruses typically can enter host cells without relying on external forces. Plant viruses can be transmitted vertically, from parent to seed. Plants are immobile, so their viruses rely on transmission for dispersal, with many plant viruses closely reliant on insect vectors [11]. As a consequence, plant viruses tend to be host generalists but vector specialists [12]. Plants lack highly specific immune systems, relying instead on generalized defenses; consequently, plant viruses can be selected to resist common mechanisms of host defense [11]. Plants can be persistently and asymptomatically infected by viruses, particularly those in the family *Partitiviridae* or the genus *Endornavirus* [13]. In comparison with crops that often display disease phenotypes, wild plants can be infected by viruses without such phenotypic indicators, leading to their prevalence being generally underestimated and underappreciated [10]. These types of infections can lead to complex, long-term dynamics over the course of a plant’s lifespan, which may vary dramatically from the population dynamics of viruses that cause acute infections.

Nearly all plant viruses have genomes composed of RNA, with most being +ssRNA [14]. This is in contrast to other well-studied viral communities—marine viromes—which are dominated by dsDNA viruses [15–20], although some data suggest that perhaps half of marine viruses have RNA genomes [21]. Due to their typically high mutation rates, RNA virus populations are expected to harbor extensive genetic variation and to be capable of rapid evolution. These properties may foster RNA virus ability to jump (emerge) easily onto new host species and to cause pandemics, relative to DNA viruses [22]; however, [11] found no difference between host range of RNA and DNA plant viruses.

We chose to focus our sampling efforts on white clover (*Trifolium repens* L.), an herbaceous perennial plant that is geographically widespread in temperate regions, ranging from the subarctic to the subtropical [23, 24]. Originally cultivated in the 16^th^ century in the Netherlands [25], this legume serves as a vital forage crop but also thrives as a weedy species in disturbed, moist areas, particularly mowed lawns. Agriculturally, *T. repens* often composes only a small proportion of a grass-legume mixed sward but provides high nutritional quality as forage. As a cover crop and companion to grass species, it is often used to enrich the soil, because it is capable of symbiotically fixing atmospheric nitrogen [26]. As an obligate outbreeding allotetraploid, cultivars and natural populations of *T. repens* are extremely diverse [24, 27, 28]. Additionally, since *T. repens* has extensive natural populations and often grows—intentionally planted or not—in and around farms, it may act as a reservoir for viruses that infect agricultural plants.

Previous studies suggest that *T. repens* is most commonly infected by the aptly-named White clover mosaic virus (WClMV). This (+)ssRNA monopartite virus (genus *Potexvirus*, family *Alphaflexiviridae*) has a 5.84 kb genome packaged in a non-enveloped, flexuous filamentous capsid. WClMV infection causes mosaic-like and mottle-like symptoms but cannot be distinguished by these discolorations alone, as its symptoms are very similar to other clover viruses. WClMV has a global distribution and a narrow host range [29, 30] able to infect other clover species as well as broad bean (*Vicia faba*) and sweet pea (*Lathyrus odoratus*). Plants inoculated with WClMV produced a third less dry material [31]. In addition to WClMV, *T. repens* is often coinfected with Clover yellow mosaic virus (ClYMV), a sister species to WClMV [32]. Both species are transmitted vertically via seed or mechanically via contact with material from infected plants [29] with grazing and mowing being particularly important for transmission between pastures [33].

Here, we compared the dynamics of RNA viral communities associated with a single host species, *T. repens*, over the small scale with the goal of testing how much a community of viruses varies in a single patch over a short time course. We collected samples of clover from 3 plots spaced every 5 m along a transect every 2 weeks for 6 weeks in total. Since we only collected tissue above ground, the viral communities are associated with the phyllosphere. Specifically, we test if the viral communities would show similar levels of species (alpha) diversity but differ in the diversity of non-dominant virus taxa. We also characterized the complexity of the quasispecies of the dominant viral species, including performing a haplotype reconstruction and tested for a signature of space or time among haplotypes. Finally, we examined the diversity of the rare species within the viral community.

## Materials and Methods

### Sampling, Extraction and Library Preparation

Clover samples were collected along a transect consisting of three plots spaced 5 m apart in the Woods Hole Ball Field and Taft’s Playground in Falmouth, MA (41°31’42.85”N, 70°40’12.59”W, Figure S1). Each sample contained at least 4 g of aboveground plant material collected in an area of one square yard and was stored in a clean, unused zip top plastic bag. Sampling equipment was surface sterilized between samples. Samples were collected every 2 weeks, allowing *T. repens* biomass to recover in between sampling events. Hereafter, we refer to the three sampled patches as A, B and C, and the time points of sampling as *t* = 0, 2, 4 and 6 weeks. Thus, the sample from patch A at time 0 is designated ‘A0’ and so on.

Plant material was pulverized using sterile ceramic mortars and pestles in 30 mL of sterile 0.85% NaCl, incubated overnight in a 50 mL Falcon tube, and then pulverized again in 10 additional ml of buffer. To eliminate plant debris, samples were centrifuged 1h at 4000 rpm, and the supernatant was transferred to a new Falcon tube and centrifuged for 20 additional minutes. The supernatant was poured through a 0.2 micron filter. The filtrate was concentrated using a Centricon Plus-70 (Millipore, Ireland) by 20-30 min centrifugation at 4000 rpm. The concentrate was re-suspended in sterile 0.1× TE buffer (pH 8.0, Fisher Waltham, MA USA) and stored at −80° C. The plant material were transferred to 1.5-mL centrifuge with 0.85% NaCl.

RNA was extracted using QIAamp Viral Extraction kit (QIAgen Hilden, Germany), treated with Turbo DNase (Ambion Foster City, CA, USA), cleaned and concentrated using RNA Clean & Concentrator™-5 (Zymo Research Irvine, CA, USA), and converted to cDNA using Ovation RNA-Seq System V2 (NuGEN San Carlos, CA, USA). Libraries were prepared using Ovation Rapid DR Multiplex System 9-16 (NuGEN San Carlos, CA, USA), and size selection performed using 1.5% Pippin Prep cassettes (Sage Science, Beverly, MA, USA). RNA, cDNA, and libraries were quantified using Quant-iT RiboGreen RNA, PicoGreen dsDNA Assay Kits (Life Technologies, Carlsbad, CA, USA), Eukaryote total RNA pico Bioanalyzer chips, and DNA 7500 Bioanalyzer chips (Agilent Technologies, Santa Clara, CA), as appropriate. Equimolar amounts of each library were mixed and run in 1.5% Pippin Prep cassettes (Sage Science). Samples were size-selected in the range of 450-650. Recovered size-selected libraries were concentrated using Ampure beads and adjusted for optimal sequencing using Quant-iT™ PicoGreen® dsDNA Assay Kit (Life Technologies, Carlsbad, CA, USA) and DNA 7500 Bioanalyzer chips (Agilent Technologies, St Clara, CA, USA). Sequencing was performed on Illumina HiSeq 2500 in the Josephine Bay Paul Center at Marine Biological Laboratory.

### Bioinformatic analyses

The paired end reads were first adapter trimmed and quality filtered with trim_galore (https://www.bioinformatics.babraham.ac.uk/projects/trim_galore/) using a quality threshold of 25 and length of 75; this was followed by additional filtering with prinseq-lite (http://prinseq.sourceforge.net) to remove low complexity sequences (dust method, threshold 7) and to remove duplicate sequences. Reads were then mapped to the genome of the barrel clover *Medicago truncatula* (GenBank BioProject PRJNA10791) to remove host reads, and also to the genome of the bacteriophage φX which is common Illumina control, using bowtie2 (http://bowtie-bio.sourceforge.net/bowtie2/index.shtml). Reads were subsequently filtered to remove ribosomal sequences using ribopicker (http://ribopicker.sourceforge.net).

Initial read level metagenomics analysis was performed by applying BLASTx to the filtered reads against a database of all RefSeq genomes using DIAMOND (https://github.com/bbuchfink/diamond), with MEGAN (http://www-ab.informatik.uni-tuebingen.de/software/megan6/) used to map BLASTx hits to the GenBank taxonomy tree.

### Metagenomics

Only filtered reads that did not map to the WClMV virus genome were used for *de novo* assembly using SPAdes (http://bioinf.spbau.ru/spades), metagenomics of the assembled contigs was performed by applying BLASTx to the contigs against a non-redundant database of all protein sequences using DIAMOND. Metagenomic pie charts were generated using KronaTools (https://github.com/marbl/Krona/wiki/KronaTools).

### Diversity measures within dominant viral species

Ultra-high coverage of the dominant viral species WClMV allowed examination of within-species genetic diversity. WClMV was subset by filtering reads that aligned to the RefSeq genome of WClMV (GenBank accession number NC_003820.1) using bwa (http://bio-bwa.sourceforge.net); an iterative mapping approach was used to call the consensus sequence of the WClMV in each sample (involving re-alignment to the initial consensus sequence, and re-calling of the consensus). The consensus sequences were then combined with all available WClMV complete genomes on GenBank, aligned using mafft (https://mafft.cbrc.jp/alignment/software/) and a maximum likelihood tree was constructed using MEGA7 with bootstrap of 100 and the GTR (G+I) substitution model. Intra-host diversity of each sample was investigated using DiversiTools (http://josephhughes.github.io/DiversiTools/) to quantify the amount of variation in each sample and determine whether sub-consensus mutations were synonymous/non-synonymous.

After WClMV was subset, diversity within the populations was examined. Mean entropy for each sample was calculated to examine the viral complexity of the populations [34]. Shannon entropy was calculated over each nucleotide site and then averaged over the entire genome to measure the amount of “disorder” in a population. Next, the degree of polymorphism was quantified at every nucleotide site across the genome, creating a mutation spectrum [34], to assess the heterogeneity of WClMV populations. Mismatch frequency of the aligned reads to the consensus sequence was determined for each sample, and each nucleotide site in the WClMV genome was binned by the proportion of reads by the mismatch frequency at that nucleotide site. Note: although conceptually similar, this mutation spectrum differs from the “spectrum of spontaneous mutations” as described in [35], which refers to characterization of the types of mutations, *e.g*. the number of transitions and transversions, *etc*.

To further examine the polymorphic diversity within the viral populations, haplotypes were constructed using QuRe (https://sourceforge.net/projects/qure/) and PredictHaplo (http://bmda.cs.unibas.ch/software.html). Due to the ultra-high depth of the WClMV samples and in order for haplotype reconstruction to complete in a reasonable time, reads aligning to the white clover mosaic virus genome were subsampled to give an even coverage of 10,000 across the WClMV genome.

## Results

### Metavirome production and contig assembly

We characterized the diversity of viral communities associated with *T. repens* from 3 plots separated by 5 m at 4 time points separated by 2 weeks with 12 samples collected in total. Sample C4 was lost in transport, so it was excluded in our analyses. For our sample preparation, we modified a protocol originally designed to isolate viruses in human stool samples for plant tissue. Our approach produced high quality results. The number of raw reads ranged between 2-24 million reads per sample. 79-99.6% reads per sample passed our quality filters, ranging from 1.6-2.1 million reads per sample. After subsequently removing reads that aligned to *M. trunculata*, the closest relative to *T. repens* on GenBank, 76-85% aligned to known viruses with 11 unique viral 11 families represented.

Assembly of metagenomic reads produced 214-34,279 contigs per sample (μ = 13927, σ = 11613). The average length of the contigs was XX. Consistent with the read alignments, the contigs are dominated the *Alphaflexiviridae* family. The remaining 161 contigs were assigned to the families *Podoviridae*, *Siphoviridae*, *Myoviridae*, *Betaflexiviridae*, *Luteoviridae*, *Potyviridae*, *Virgaviridae*, *Closteroviridae*, *Cystoviridae*, and *Rhabdoviridae*.

### Taxonomic composition of the viral community associated with *T. repens*

We predicted that the viral communities associated with *T. repens* would be dominated by a few viral species with many more rare species. Specifically, we predicted that we would detect WClMV and ClYMV based on previous studies of viruses typically detected infecting *T. repens*. Our results confirmed that the communities were consistently dominated by these two viral species, with WClMV as the most frequent. Notably, these two species were extremely common relative to the other virus species detected in the viral community. After filtering, WClMV accounted for 50-83% of the reads in each sample, and that for ClYMV was 2.2-24%. These two species consistently dominated the viral communities, accounting for 76-85% of the filtered reads (Figure 1). Furthermore, when ClYMV accounted for only a small proportion of the reads, WClMV compensated by comprising a larger proportion of the remaining viral community rather than another viral species. We examined the distribution of the viral species within the communities associated with *T. repens* after removing WClMV and ClYMV. We found that removal of WClMV and ClYMV showed that other members of the *Alphaflexiviridae* family still dominated the remaining identifiable species (Figure 2, Table S2). 76-85% of the filtered reads aligned to known sequences in RefSeq included other plant viruses, e.g. *Clover yellow vein virus, Potato yellow dwarf nucleorhabdovirus*, and *Beet yellow stunt virus*, as well as bacteriophages (e.g. Pseudomonas phage φ2954, Klebsiella phage SopranoGao, and Shigella phage Sf11), and viruses associated with invertebrates (e.g. Hubei picorna-like virus 51 [36]).

**Figure 1.**
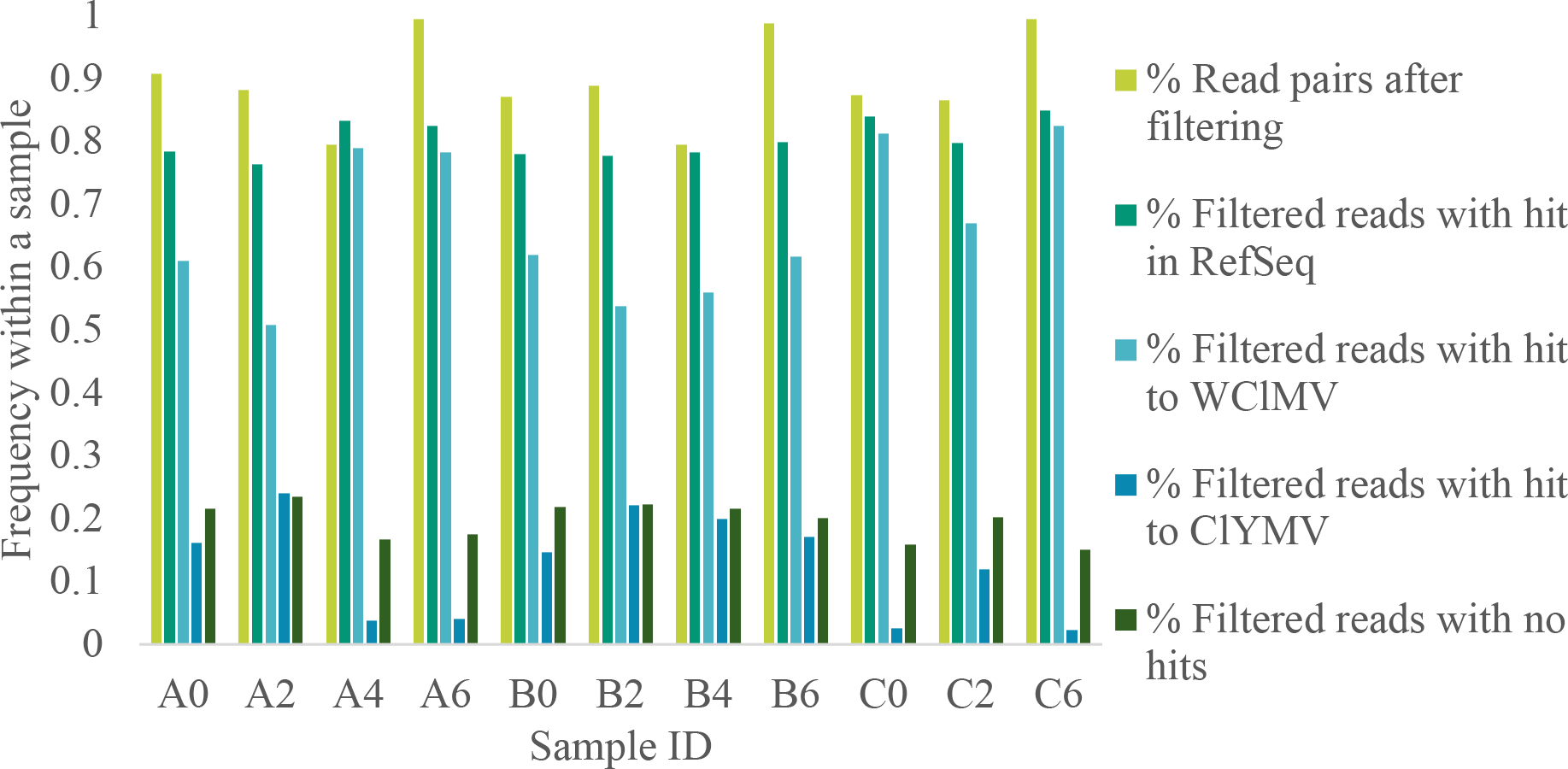
Summary of the read statistics for each sample. Plots A, B, and C were separated by 5 m with samples taken at 0, 2, 4, and 6 weeks. Bars represent the frequency of the read pairs that remained after filtering, matched an existing sequence in the RefSeq database, matched WClMV, ClYMV, or failed to match to any sequence in RefSeq.

**Figure 2.**
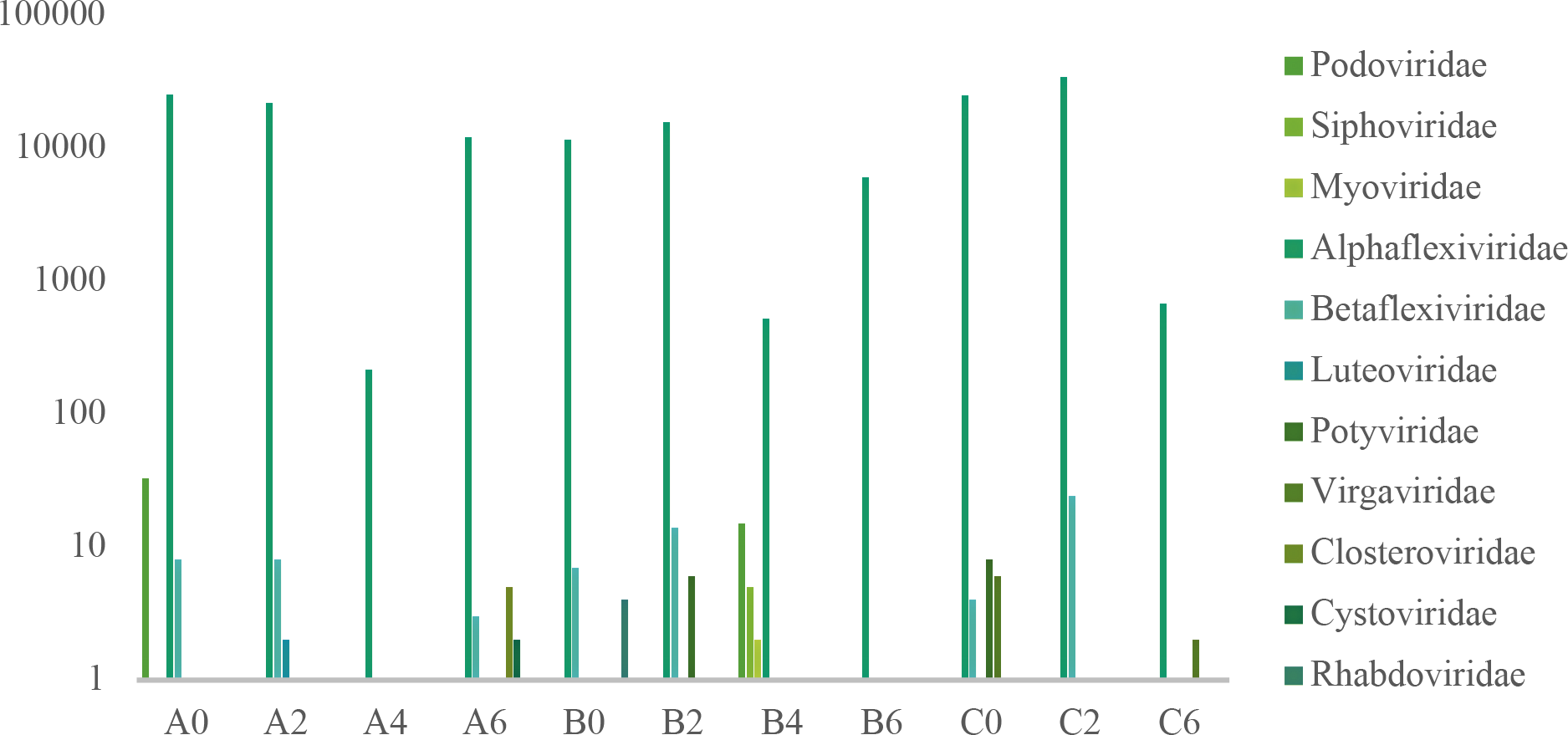
The rare virus families present in each of the samples after the dominant viral species (WClMV and ClYMV in the *Alphaflexiviridae* family) were removed. Note that the *Alphaflexiviridae* present in this sample are comprised viral species other than WClMV and ClYMV.

### Analysis of the dominant viral species: WClMV

WClMV consistently dominated the viral communities across all samples, composing approximately two thirds of all RefSeq hits (μ = 0.667, σ = 0.117). Given the high coverage (μ = 8.32 × 10^6^, σ = 4.36 × 10^6^ read pairs per sample), we were able to reconstruct the full viral genome from each sample and also analyze the dynamics of WClMV. We compared the consensus sequence of WClMV from each sample to all other complete WClMV genome sequences on GenBank on a phylogenetic tree (Figure 3). Our samples clustered with the only other sample of WClMV from North America and the original sample of WClMV sequenced from New Zealand. The other sample from New Zealand clustered with samples from South Korea and Japan. As for our samples, all time points from plot B clustered together as did the majority of samples from plot A; sample A2 is an outlier that does not cluster with the other A samples. Although plot C samples do not form a distinct clade, they cluster with the other WClMV consensus sequences from this study.

**Figure 3.**
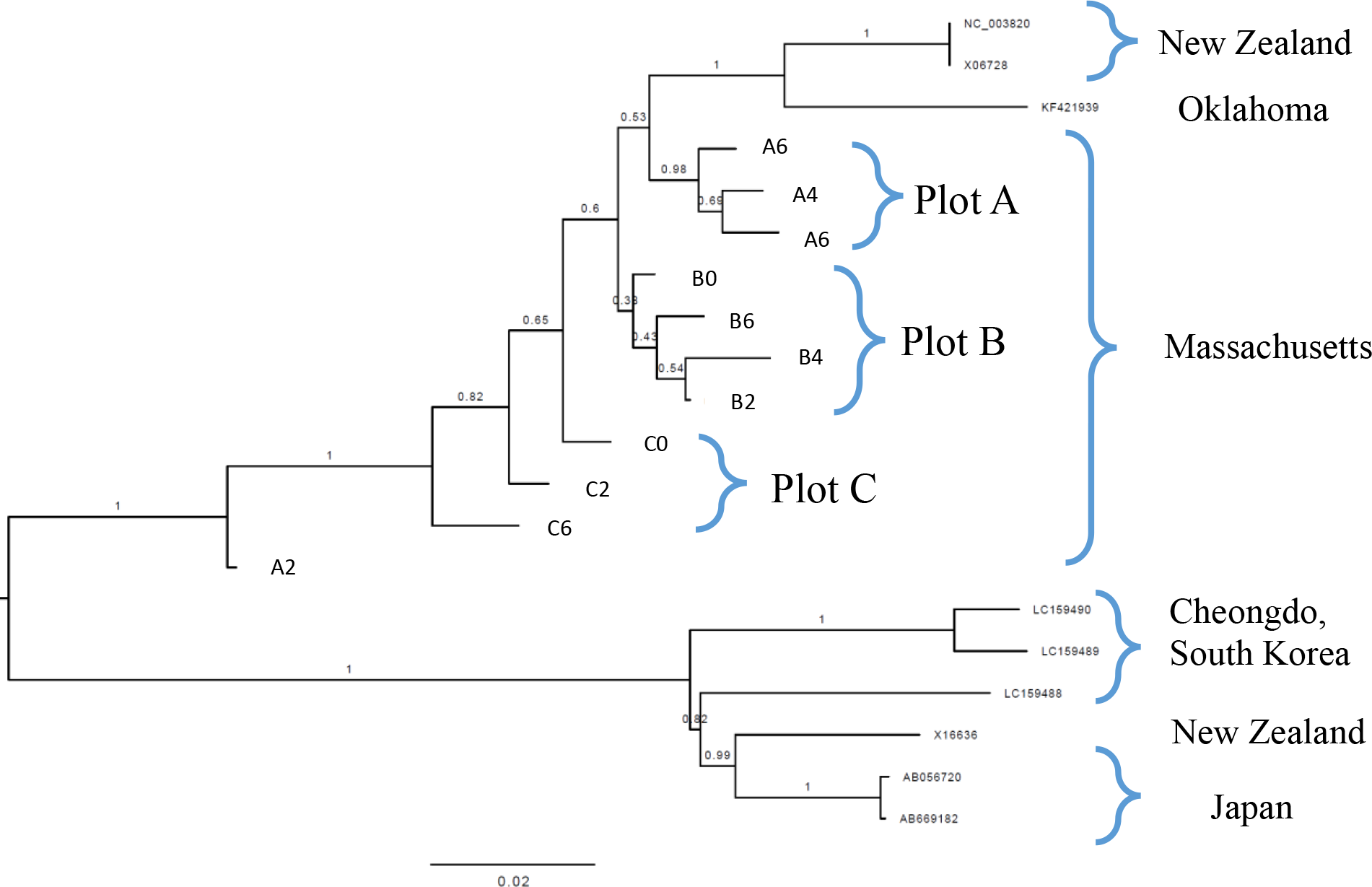
Phylogenetic tree for all samples of WClMV combined with all complete genome sequences of WClMV from GenBank.

In order to examine the dynamics of WClMV, we characterized the complexity of the viral quasispecies within each sample using Shannon entropy. This metric assesses the diversity of the genomes within a sample. The mean Shannon entropy per nucleotide site within a sample was relatively high with values between 0.105-0.109 for all time points at plots A and B. However, the entropy varied more for plot C ranging 0.093-0.106 (Figure 4A). Of the 5843 nucleotide sites, the number of nucleotide sites that is invariant (Shannon entropy = 0) is 1-103 per sample (μ = 25.5, σ = 31.7), although this may be due in part to the high coverage. Next, we generated a mutation spectrum [37] to examine the frequency of mismatches [non-reference bases] across all genome positions (Figure 4B). The spectrum groups genome positions by their total mismatch frequency, to give an overview of the average mismatch frequency throughout the genome. We compared the mutation spectrum of the samples against the spectrum of another RNA virus, namely foot-and-mouth disease virus (FMDV) from a single foot lesion [38] for reference. The samples of WClMV clearly have a substantially higher heterogeneity than the FMDV sample, with a large proportion of genome positions displaying mismatch frequencies greater than 10%. Further investigation of these mutations showed that the vast majority were non-synonymous suggesting that these were functional mutations and not mutation errors introduced during sequencing or sample preparation. Due to the number of high frequency mutations observed within the viral population, we suspected that the sample contained multiple strains of WClMV and therefore proceeded to haplotype reconstruction. The phylogeny of the reconstructed haplotypes (Figure 4C) suggested that the haplotypes could be divided into two different groups with high confidence, supporting the hypothesis that two different strains of WClMV coexisted within the samples.

**Figure 4.**
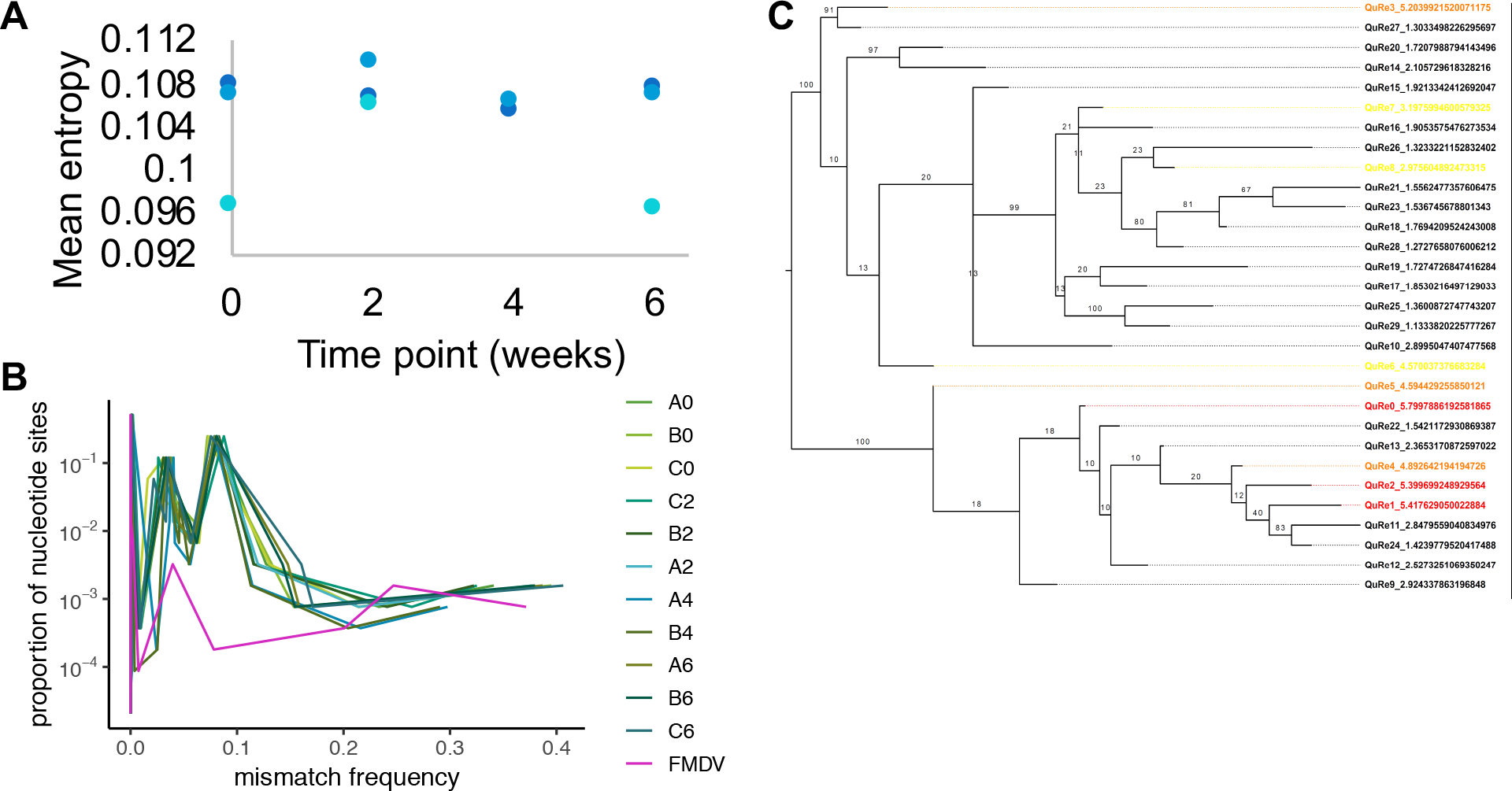
Shannon entropy of the WClMV of each of the samples. A) Shannon entropy (averaged over all genome positions for each sample) for each plot over time. Samples collected from plot A are in green, B in light green, and C in chartreuse. B) Mutation spectra of each sample. The mutation spectrum for FMDV (Orton et al 2013) is included as a reference. C) Haplotype reconstruction of WClMV pooled across sample.

## Discussion

We sampled the RNA viral communities associated with a single host species over a scale of meters and weeks. We found a striking pattern of viral diversity in the communities associated with *T. repens*: all communities were dominated by a single viral species, WClMV. The second most abundant species was its sister species, ClYMV. Most of the remaining rare species were also members of the *Alphaflexiviridae* family, but there were no discernible patterns of diversity among the very rare viruses outside this family. Because we were able to recover a high level of reads per sample, we were able to identify diversity within a species rather than higher taxonomic ranks. Although we saw low diversity at the community level, we observed very high levels of diversity within the dominant species. This enabled us to examine the population dynamics of WClMV further. We found more similarity among WClMV reads by sampling location rather than time point and the consensus sequence for each of our samples tended to cluster together when compared with all other whole genome sequences on GenBank. Also noteworthy is that such a high proportion of the filtered reads (76-85%) aligned with known viral genomes on RefSeq. For instance, only 10% of the virus-derived sequences from plants belong to known virus species [39]. This may be explained by the fact that our samples are dominated by known viruses.

Since we were able to recover such a high number of reads per sample, we want to highlight the steps in our sample preparation that we believe contributed to our success. We modified the sample preparation as described by Minot and colleagues [40], which isolated DNA viruses from human stool samples, for plant samples. Similarly, we used a large plant sample, ground it, and mixed it with a large volume of buffer. The large volume allowed us to mechanically lyse cells and separate cellular debris from the plant while the filtration steps allowed us recover viruses in a concentrated form. Unlike Minot et al [40], we opted to omit their chloroform step to avoid disrupting membrane-enclosed viruses. Additionally, they used a different set of kits for genome extraction and library preparation, because their study focused on DNA viruses. Regardless of those differences, both studies recovered a high number of reads (ours: 2-24+ million reads, theirs: 15-39 million) per sample. To our knowledge, this is the first publication of this methodology as applied plant viral metagenomes. This approach ultimately proved successful at separating viral and host genomes: we found that >10% of the samples aligned with the host reference genome, *Medicago trunculata*, a relative of *T. repens*. In comparison with other methods for isolating viral communities associated with plants, this method is relatively unbiased (see [39] for a review). The only amplification step occurred during the library preparation; the kit amplified samples with random primers so no bias is anticipated. Overall, this approach is a promising method for examining viral communities associated with plants.

The high diversity that our populations exhibited suggest that these plants may have long-term WClMV infections in *T. repens*. WClMV is a long-term infection, rather than an acute infection, allowing viral diversity to arise and accumulate via mutation. Since *T. repens* is a perennial plant and can survive for many years, these viral infections may also be several years old. Additionally, these samples were collected from a municipal ball field that is regularly used, meaning that normal use and mowing can allow mixing of viruses from throughout the field. Furthermore, the mowing may introduce virus from other recently mown fields. These factors may help account for the high Shannon entropy values. In comparison, the Shannon entropy values for individuals experimentally inoculated with an acute infection of Foot and Mouth Disease Virus (FMDV) maxed out at 0.0008 [34], while the values for our populations peaked at 0.11.

The second most abundant viral species in our communities was ClYMV. This virus has also been associated with white clover for decades. Interestingly, ClYMV was historically limited to the Western US [30]. However, a more recent a disease note documented ClYMV east of the Mississippi River for the first time, in *Verbena canadensis*, which is in a different family from *T. repens*, in Florida [41]. Furthermore, ClYMV appears to be expanding its geographical ranges recent reports for the first time in the UK [42] and in *T. repens* in Australia [43]. Taken together, this suggests that ClYMV may be expanding both its host and geographical range, first jumping into ornamental *Verbena* spp. and then moving globally through the sale of live plant material [44].

Beyond the two dominant viral species, most of the viruses we detected were also from the family *Alphaflexiviridae*, but we were not able to identify many of these reads beyond the XX taxonomic rank. Although we were not targeting DNA viruses with our methods, we did detect several DNA viruses present among the rare species, which we likely detected during transcription. Specifically, we detected three tailed phage families—*Myoviridae*, *Siphoviridae*, and *Podoviridae*—all of which have a linear dsDNA genomes. This is consistent with other studies, *e.g*. the findings from a study of viral communities associated with *Capsicum annuum*: despite extracting RNA from the plant tissue, ~3/4 of the viral reads aligned DNA viruses [45]. If these results are real, this suggests that this method was able to detect DNA viruses during transcription and that these phages were actively replicating during infection of bacteria associated with the plant.

## Acknowledgements

We thank researchers in the Josephine Bay Paul Center at Marine Biological Laboratory for helpful advice and useful comments on early versions of the manuscript.

